# Machine Learning-based state-of-the-art methods for the classification of RNA-Seq data

**DOI:** 10.1101/120592

**Authors:** Almas Jabeen, Nadeem Ahmad, Khalid Raza

**Affiliations:** Department of Biosciences, Jamia Millia Islamia, New Delhi-110025, India; Department of Computer Science, Jamia Millia Islamia, New Delhi-110025, India

**Keywords:** RNA-Seq data, Machine Learning, Deep learning, Classification, Clustering, Feature Selection

## Abstract

RNA-Seq measures expression levels of several transcripts simultaneously. The identified reads can be gene, exon, or other region of interest. Various computational tools have been developed for studying pathogen or virus from RNA-Seq data by classifying them according to the attributes in several predefined classes, but still computational tools and approaches to analyze complex datasets are still lacking. The development of classification models is highly recommended for disease diagnosis and classification, disease monitoring at molecular level as well as researching for potential disease biomarkers. In this chapter, we are going to discuss various machine learning approaches for RNA-Seq data classification and their implementation. Advancements in bioinformatics, along with developments in machine learning based classification, would provide powerful toolboxes for classifying transcriptome information available through RNA-Seq data.

## 1. Introduction

In day to day life, we are verge to encounter various infectious agents like virus, prion, fungi, bacterium, etc. To a great extent our body provides primary defence mechanism to the infections caused by them but not necessarily all. The emergence of superbugs (antibiotic resistance bacteria) such as *Staphylococcus aureus* (MRSA), *Klebsiella pneumonia* (CRKP), etc and recurrence of influenza, zika, ebola, etc, are of great concern in today’s world and even in future. Studying these infections and diseases may find a great importance in upcoming years. With the advent of digitization and availability of high-throughput modern devices at cheaper rates, the task of sequencing has been boosted up resulting in massive generation of heterogeneous data (Big Data) at lower experimental and computing cost. Recent advances in the field of sequencing, i.e., Next Generation Sequencing (NGS) via RNA-Sequencing (RNA-Seq) method enabled the biologists and biotechnologists to measure the expression levels of several transcripts simultaneously. By applying such information one is able to develop various classification algorithms based on expression levels. It is now considered an emerging method for disease classification and diagnosis and identification of potential biomarkers of disease [2]. They are also being used to identify exogenous RNA contents (Viruses) from RNA-Seq data. The information from RNA-Seq data utilized for this were earlier overlooked but by employing machine learning methods through classification algorithms, these features can be easily identified in RNA-Seq data. It is noteworthy that, out of several only few information from RNA-Seq data is useful for identification hence prediction which are considered as Target rich data that are needed to be inferred. For this reason machine learning techniques find a great usage in bioinformatics to perform the real time predictive and analytical study in order to give rise to intelligent informed decision [1].

It is observed that two similar disease state samples which differ categorically may not able to be differentiated accurately in presence of background noises that matches. Thus it is necessary to select the key differences via machine learning techniques [3]. “Classification” has been a great topic for research in fields of machine learning in recent years as it has found a great applicability which searching for disease biomarker and drug targets [4]. Various computational tools have been developed for studying pathogen or virus from RNA-Seq data by classifying them according to the attributes in several pre-defined classes, but still these tools and approaches to analyze complex datasets are still lacking. Moreover there is no ‘one fit to all’ technique for classification and analysis of RNA-Seq data which makes disease classification even more challenging. In addition, the high dimensionality incurred in NGS data, including RNA-Seq, has thrown various challenges including curse of dimensionality problem, and hence several existing classifiers cannot be directly applied.

In this context, there is an immediate need of upgrading current classification approaches to meet the analysis of high-dimensional data such as RNA-Seq data. In this chapter, various machine learning approaches for RNA-Seq data classification are discussed with their pros and cons. The development of various classification models is an emerging area of research for disease diagnosis and classification, examining it at molecular level as well as discovering its potential markers which could be targeted in disease identification and drug discovery. These developments would further allow transcriptomic analysis in rare cell types and cell states, and also would enable reconstruction of biological networks at cellular level. These bioinformatics advances, along with developments in machine learning based classification, would provide powerful toolboxes for classifying transcriptome information available through RNA-Seq data.

## 2. RNA-Seq and its analysis pipeline

RNA-Seq is found to be a powerful transcriptome profiling technique of NGS which can provide a detailed view of RNA transcripts in test sample. RNA-Seq measures expression levels of several transcripts simultaneously [72]. The general methodology for RNA-Seq analysis concerning a disease involves expression analysis from raw sequence data of disease after normalization, then constructing individual network modules for gene identification followed by examining their aggregate network properties [5]. This system biology approach can successfully identify multiple genes related to a disease. These identified genes can then serve as target for further drug discovery process. Fig.1 represents the systematic steps for RNA-Seq analysis method [6].

**Fig. 1.**
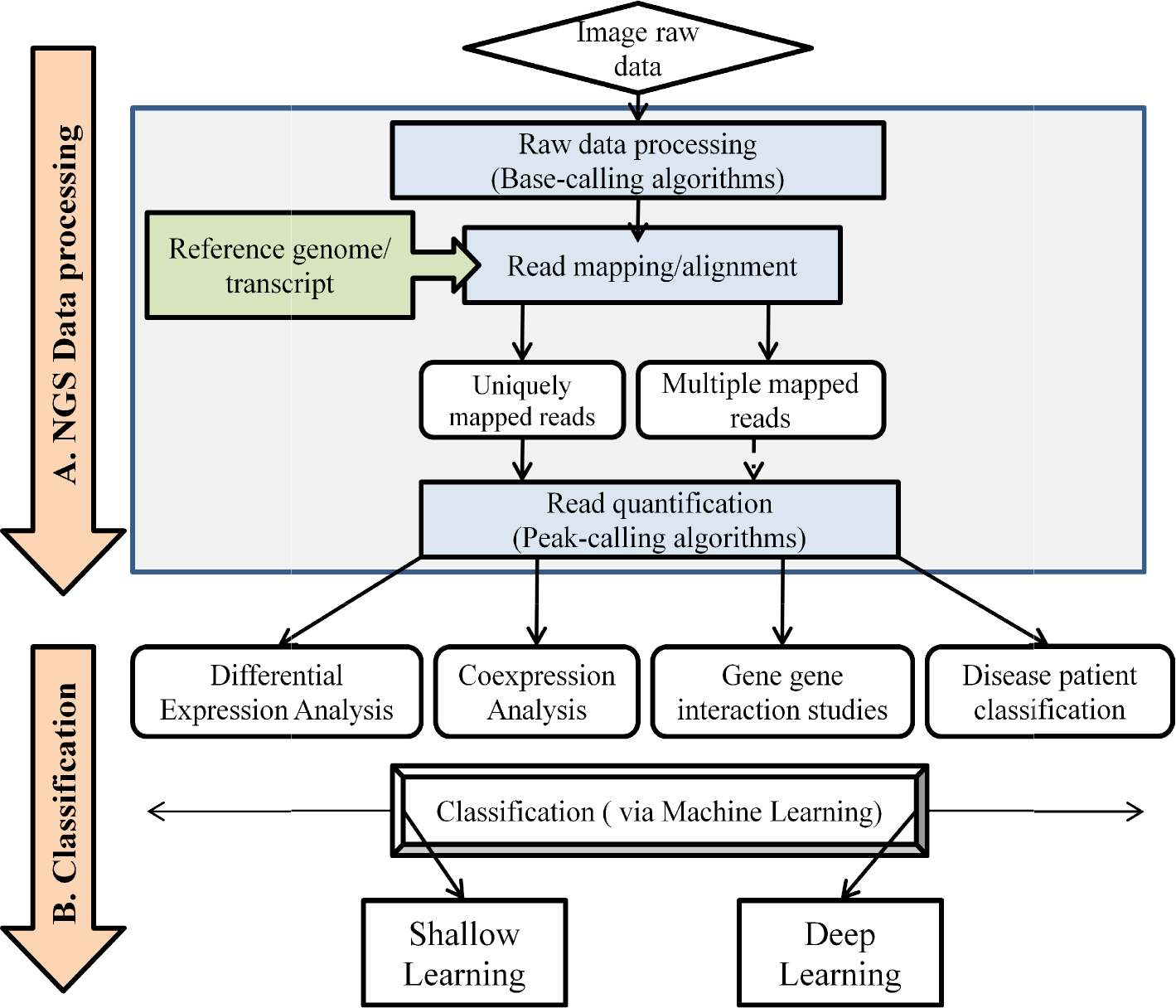
Steps involved in RNA-Seq data analysis pipeline. A. The NGS data (RNA-Seq dataset) is preprocessed and transcripts get quantified which are subject for differential expression analysis, co-expression analysis, gene-gene interaction study or disease patient classification. B. The result undergoes classification procedure to classify elements according to their attributes by machine learning algorithms [6].

The key hub genes associated to the disease pathways obtained from differential expression analysis undergoes for further classification. The reads identified through this process can be an exon or other region of interest. [1]. Machine learning is an art of learning that doesn’t involve any sort of explicit programming. It can occur in either of the two forms: conventional “Shallow” learning or “Deep” learning [7]. The typical machine learning workflow for classifying of data from computational point of view can be seen in Fig. 2 [63]. Similar approach can be implemented when dealing with RNA-Seq data.

**Fig. 2.**
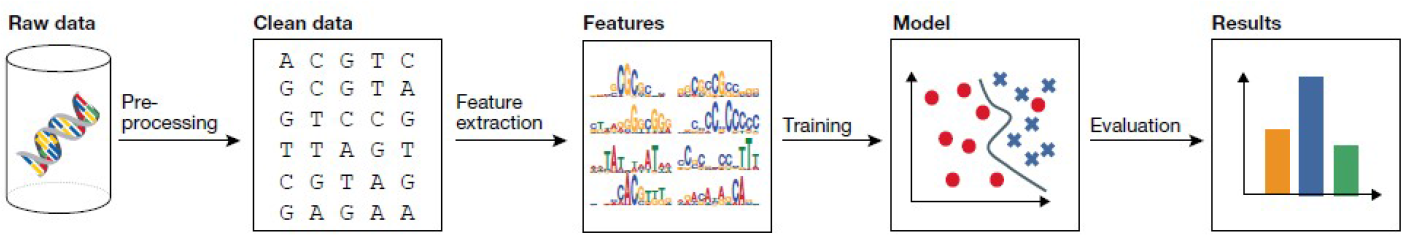
The classical machine learning workflow consists of four steps: data preprocessing, feature extraction, model learning and model evaluation [63].

## 3. Feature selection via data pre-processing

### 3.1. Features involved in RNA-Seq Classification

Feature identification is the most important step in construction of a machine learning classifier. Feature sources such as size of RNA transcript or GC content is not enough to build an efficient discriminating identifier for RNA transcripts. Other features such as graph features from sequence, conservation score features, component composition features, ALU repeats and tandem repeats, and ORF features for transcript sequences can also be a good feature sources for training and testing a model [8].

**Table 1.** Features for microRNA detection and analysis [13].

| S. No. | Feature | Description |
| --- | --- | --- |
| 1. | Read count | Number of reads mapped to pre miRNA |
| 2. | Length | Length of longest hairpin structure |
| 3. | MFE | Mean free energy of hairpin |
| 4. | GC | GC content of small hairpin |
| 5. | Loop GC | GC content of the loop |
| 6. | Asymetric bulges | Number of Asymetric bulges and mismatches regarding the stem |
| 7. | Symetric bulges | Number of Symetric bulges and mismatches regarding the stem |
| 8. | Bulges | Number of bulges in stem |
| 9. | Longest bulge | Number of non-pairing nucleotides of the longest bulge |
| 10. | Mismatches pre-miRNA | Number of single mismatch in the hairpin |
| 11. | Mismatch miRNA | Number of single mismatch in the mature miRNA region of hairpin |
| 12. | Stability | Frequency of original hairpin structure found in the elongated structure. |
| 13. | Alternating stability | Reports that structure disappears when extending the sequence, but reappears again. |
| 14. | Triplet SVM features | All features that were proposed by Fan & Zhang, 2015 [14]. |

### 3.2. Steps for classification model building

The machine learning methods for RNA-Seq data classification require pre-processing of datasets in order to raise chances of getting effective results. The necessary steps for a typical classification process are outlined in Fig. 3. It comprise of basically three steps as Feature selection followed by classification model building and then validation of the constructed model.

**Fig. 3.**
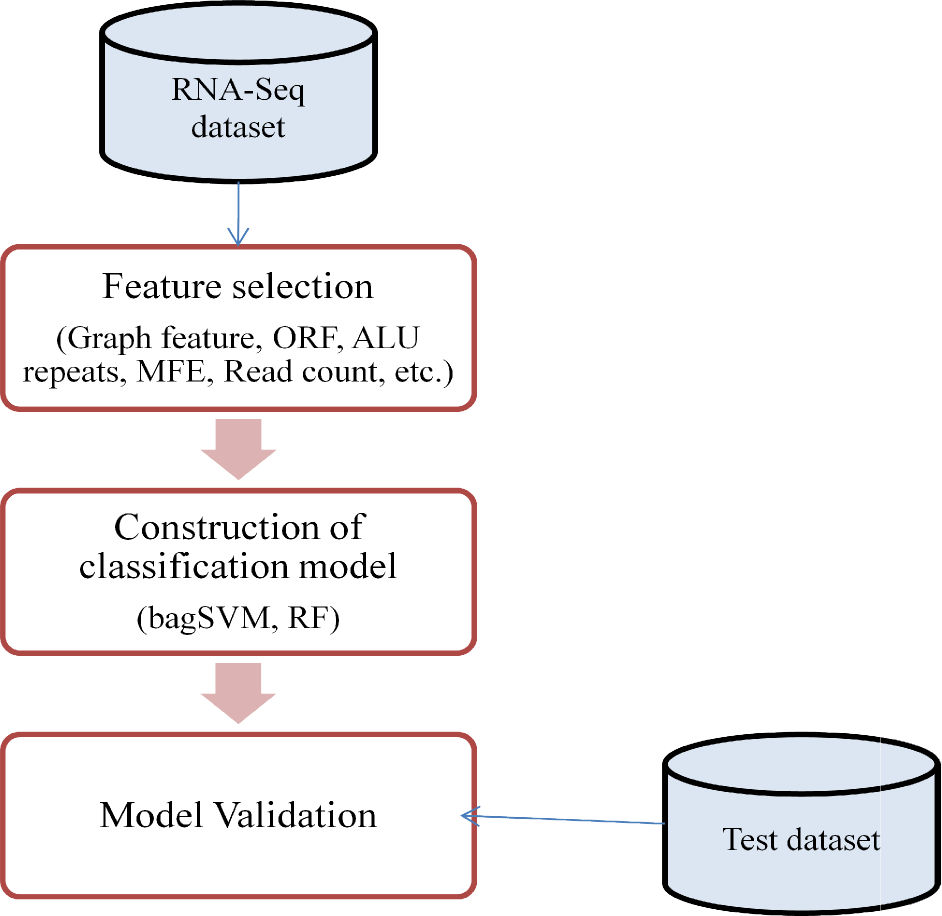
Steps involved in a typical Classification process.

In feature selection step, it is necessary to create optimal data subset which reduces the noise and biases hence alleviate accuracy of classification process. It also reduces the effort and cost for computational aspect. The optimal data subset, thus consist of a more interpretable feature. The classification model building step deals with constructing classification models by implementing various machine learning algorithms. With help of the selected features, machine learning algorithms learn the classifiers with definite parameters from the training dataset. Thus the built model can predict the assignment of objects (biological sample) to a specific class. A typical feature selection involving Random forest classifier is shown in Fig. 4 [42].

**Fig. 4.**
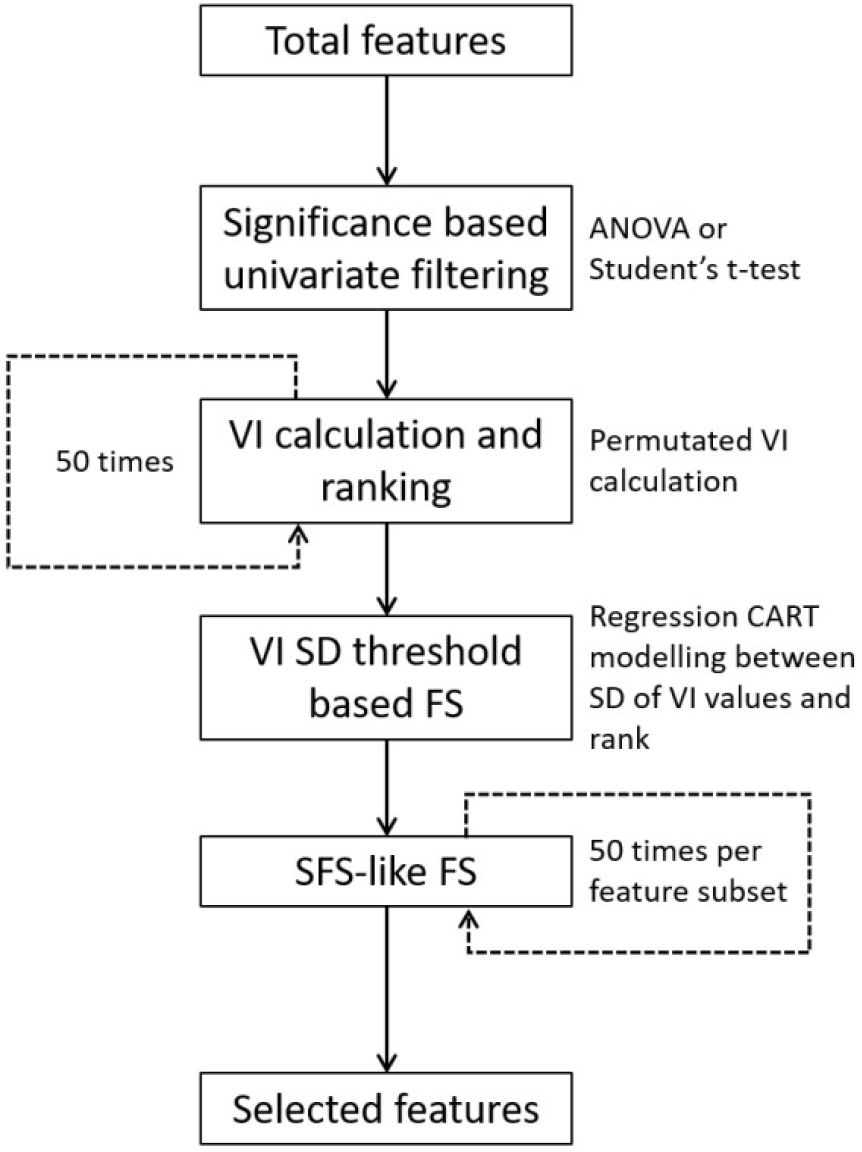
The flowchart of the current RF-FS implementation [42]

The commonly used classifiers in model building process includes Support Vector Machines (SVM), Bagging SVM (bagSVM), Random Forest (RF), CART, Linear discriminate analysis (LDA), Artificial neural networks (ANNs) and k-nearest neighbours. Model validation step is critical step to cross check the efficiency of the model. Various model validation approaches like Holdout validation, k-fold cross-validation and bootstrapping have been emerged till date [2].

### 3.3. Methods for Feature selection process

Feature selection process is used to extract the relevant features to be implemented in classification model and remove the unimportant, redundant features in order to reduce the curse of dimensionality. This would make the learning process for classification time efficient and will increase the performance of model [15]. For a Big Data feature selection process such as in case of RNA-Seq data, both supervised learning and unsupervised learning can be implemented to make decision. Furthermore, ranking of the features according to their relevance to the classification problem and then selecting the best ones out of them can improve the performance of prediction model [16].

Expression level: The basis of disease diagnosis via classifying data into different classes is simply measuring the changes in expression level across the classes. Several studies adopting this approach have been emerged to determine important biomarkers (features) in order to predict survival outcomes, disease subtypes, drug sensitivity and even behavioural characteristics [17, 18, 19, 20].

#### Differential Expression (DE) based classifiers

The difference in expression level (Differential Expression (DE)) of genes is found useful in classification in order to identify disease biomarker [20]. For RNA-seq data, classification is done by edgeR package [21] where the genes are ranked on basis of their likelihood ratio in test statistic result obtained from negative binomial linear models. After transformation of RNA-Seq count data in order to eliminate overdispersion, the sample dataset is subjected to the Poisson linear discriminant analysis (PLDA) machine learning approach to efficiently find the correct decision boundary and do predictions at high accuracy [22]. It was observed that expression variability (DV) classifier which was based on adaptive index models [23] outperformed a differential methylation classifier in context of early-stage cancer prediction. This suggests that traditional differential expression (DE) classifiers ignore important differences which are present in real data sets [20].

#### Expression Variability (DV) based classifiers

When deregulation in process signalling occurs, it leads to change in expression variability (DV) of target genes. DV exploits the characteristics of deregulated networks to identify and assess important biomarker for an improved disease prediction and treatment. But, it also ignores some useful information from changes in locations between classes [20]. For RNA-seq data, logarithmic transformation was done by implementing DESeq2 just to avoid mean-variance based traditional transformation approach, hence also avoided the expression variability caused merely by differential expression [24]. Features then ranked then selection is applied. Prior to the training and prediction step, the value for each feature based on difference in each measurement with median of all samples in a training dataset. Then Fisher’s linear discriminant analysis (FLDA) method can be employed over them for classification [20, 25].

#### Differential Distribution (DD) based classifiers

Biologically, a change in distribution (unimodality to multimodality) suggests the expression range of genes for a normal cellular functioning. A novel kernel density-based DD measure with a prognostic algorithm and showed that it performs well in terms of classification and stability on both simulated and three sets of real high-dimensional transcriptome data. DD classifiers are advanced classifiers which simultaneously identify Differentially Expressed Genes (DEGs) and also Differentially Variable Genes (DVGs) or both. DD aims to avoid the need for ad-hoc DE and DV classifier aggregation algorithms [20].

For the RNA-seq data set, firstly the counts should be log transformed to prevent biasing of feature selection towards differentially expressed genes due to overdispersion of count data. Naive Bayes classifier selected each feature to the sample dataset. The DD classification using kernel density estimate voting showed a better performance than LDA [26]. This helps to conclude that DD metrics can be better discriminative measure for genes identification. Moreover this DD metric can be efficiently used in a novel classification scheme of RNA-Seq datasets.

Assessment of genes via classification through DV is desirable when sample with experimentally unknown genes are required to be classified. But due to lack of availability of variability-associated disease genes in public database, only DV and DD [27] are taken into account for classification. A graphical summary of DE, DV and DD features selection method with classifiers are shown in Fig. 5 [20].

**Fig. 5.**
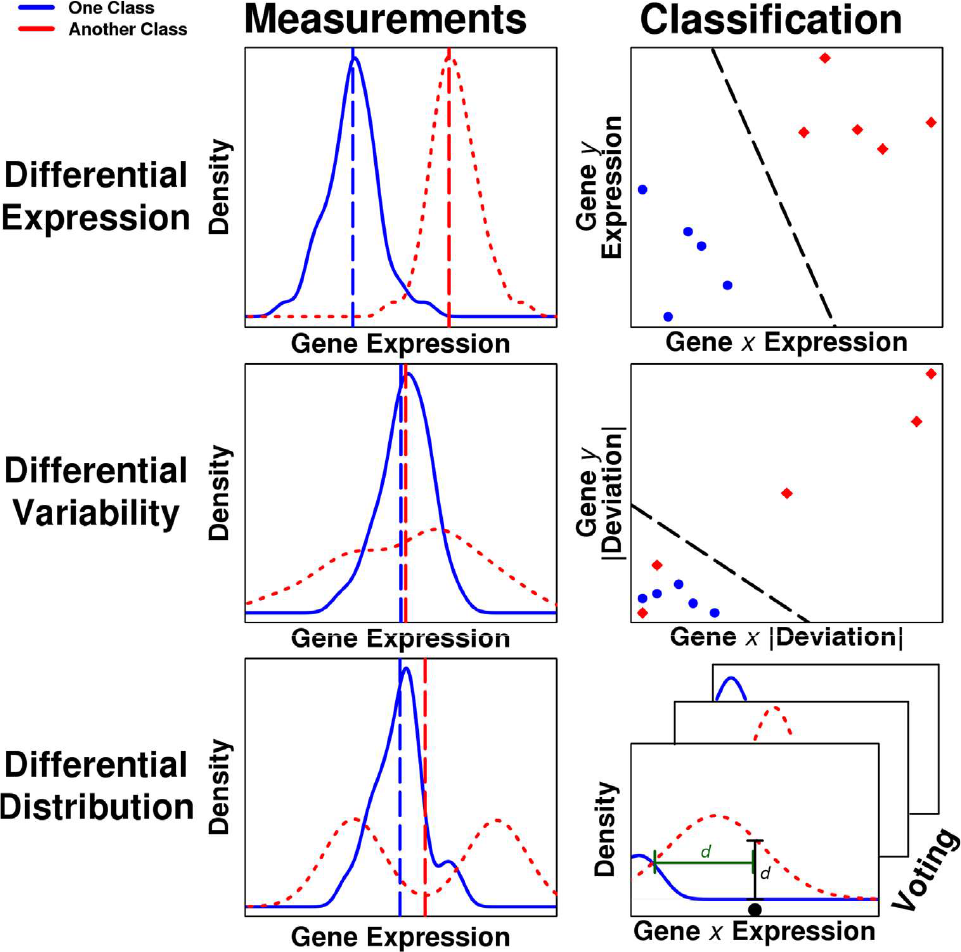
Summary of the feature types and classifiers. For each of differential expression, differential variability and differential distribution, a representative gene profile is given and an illustration of the classification process given. In the left column, the dashed vertical lines represent the means of the class distributions. In the right column, the variables x and y denote two different genes in a data set. The bottom right panel illustrates that each gene from the selected gene set votes independently in differential distribution classification [20].

It is observed in [20] that the performance of all three classification schemes based on their prognostic error rate and biological relevance, the DV classification has inefficient feature selection but good error rate under simulation. Differential distribution selection identifies different sets of disease related. Differential distribution is the most stable method of ranking and selecting features.DE classification can only detect changes in means of datasets and can possibly neglect signatures associated with transcriptional deregulation.DD classification is, therefore the better approach for gene expression assessment with good classification accuracy features than DE or DV selection for biologists to start an experiment [20].

## 4. Machine learning approaches for classification of RNA-Seq data

Machine learning is the field of computer science that involves efforts in development of various computational methods that learn from training data. It is divided into two approaches Shallow learning and Deep learning. Shallow learning consists of neural networks with single hidden layer or SVMs. They are simply supervised and unsupervised learning methods [28]. The supervised learning methods rely on classifiers whereas unsupervised learning implements clustering algorithm. In a supervised learning model learns from a set of predefined objects with class label (training set). The knowledge inferred from it used to classify the unknown objects (test objects) accordingly. Whereas, unsupervised learning do not depend on the availability of prior knowledge (training data sets) with class labels. Deep learning consists of neural network with several hierarchical layers. It is a good alternative for big data analytics with high accuracy. The traditional machine learning methods are found inadequate in handling voluminous data using the current computational resources Therefore Deep learning evolved but still further development is needed. The categorization of machine learning can be seen in Fig. 6.

**Fig. 6.**
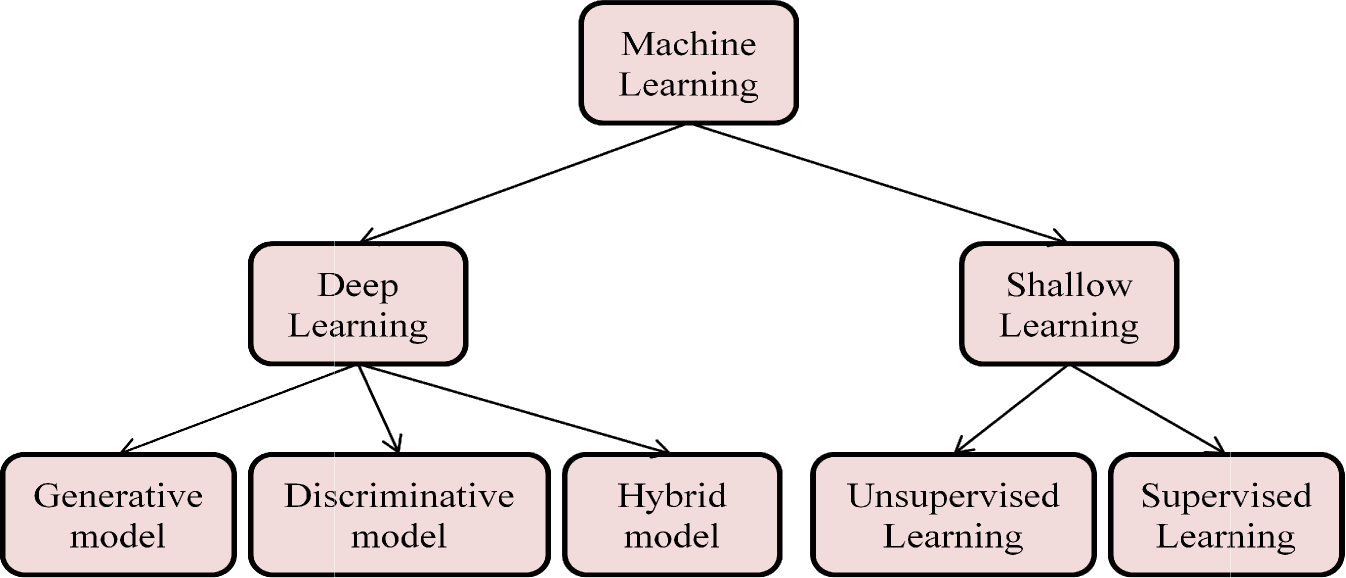
Categorization of machine learning technique.

### 4.1 Supervised learning methods

In a supervised learning, labeled training datasets trains the supervised learning model to predict the class label of test objects based on knowledge inferred from training dataset. Supervised learning methods can further be subclassified into classification models (predicting categorical outputs) or regression models (predicting continuous outputs) [1] as in Fig. 7.

**Fig. 7.**
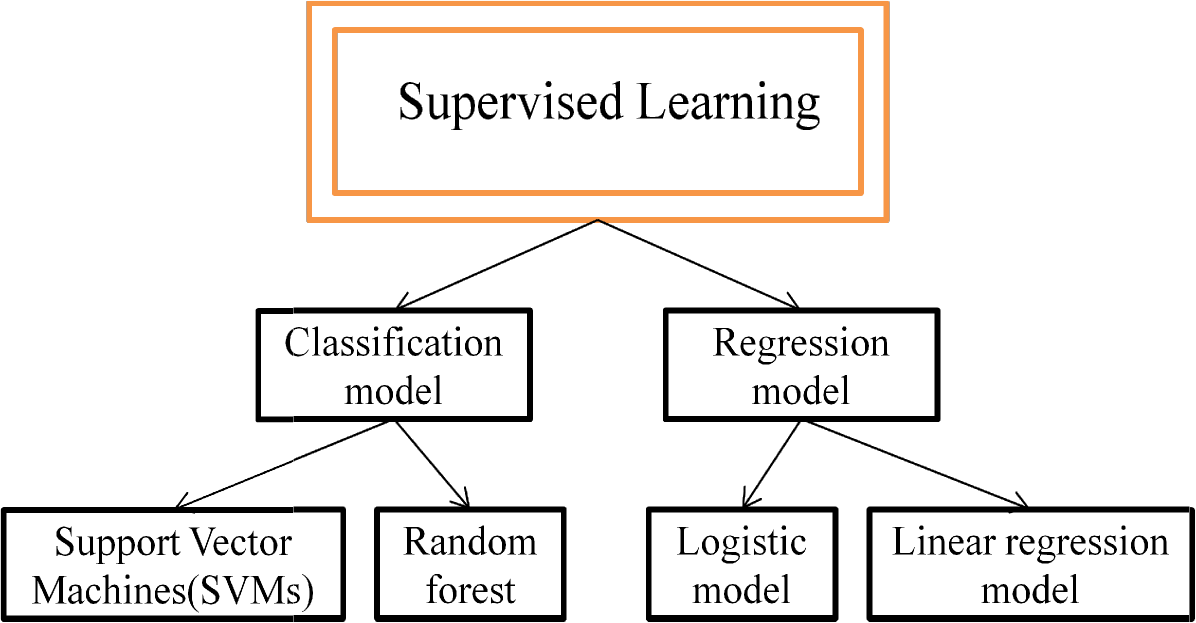
Categorization of Supervised learning process.

#### Ensemble learning theory

Machine learning methods are widely used in research, data mining, medical diagnosis, etc. One of the approach called Ensemble learning” is that branch of machine learning that applies ‘4 eyes see more than 2’ ideology. It learns a model implementing several weak learners, using specific rules, than integrates the learning result from each to lay a final efficient, predictive result. Therefore, ensemble learning yields more efficient machine learning than a single learner [4]. Ensemble method for learning improves the predictive performance of a classifier in generating an accurate model. It is technically a collection of several base (weak) classifiers whose individual decisive result is summed up in a way that can classify new data points more effectively [2, 29, 30]. Fig. 8 shows the basic concept behind ensemble learning [4].

**Fig. 8.**
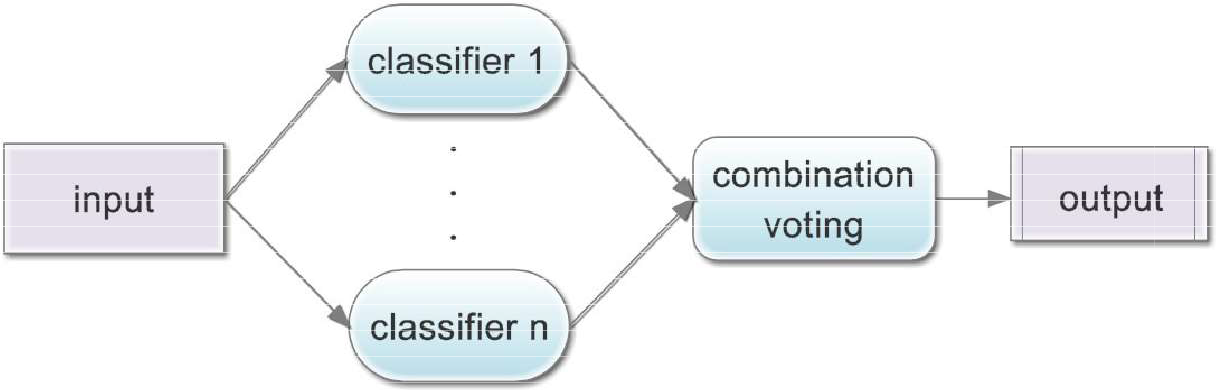
Basis of a typical Ensemble classification [4]

At the very first, a study on boosting model [31] concluded that certain combinations of weak classifiers (base classifiers) have same performance as strong classifiers, thus concluding that identification of a strong classifier is not needed. Reasons for ensemble learning approach to be more promising than traditional single classifier based learning can be summarized as follows[29, 32]:

- Due to limited number of the training sets used in single classifier learning, a learning algorithm is unable to precisely learn to target. But this is not so in case of ensemble learning and thus, here the offsetting error between assumptions and extracting the classification by integrated classifiers proven to be advantageous statistically
- The machine learning analysis implementing artificial neural networks and decision trees proven to work better for learning hypothesis for a non-deterministic polynomial dataset, as it can easily be incorporated in other classifiers. This gives an idea that when several of such assumptions are combined together will give a synergistic upraise to classification result to actual target function value and thus, ensemble will proven to be better computationally too [4].
- Stability of feature selection process is necessary while resampling procedure in order to ensure that selection was not merely a coincidence. Ensemble feature selection method ensures the feature stability [33].

Ensemble approach though is an effective and accurate classification approach but still is computationally costly as it requires the training of many similar models. Moreover, bagging up the features from different single classifier models depends on user-specified parameter [4]. A large number of models include single classifier such as linear and nonlinear density-based classifiers, K-nearest neighbor (KNN), decision trees, naïve Bayes, SVMs, neural networks and the others are the ensemble of several single classifiers such as Random Forest, bagSVM, etc. It can be concluded that for an efficient classification solution one must combine experts from various specific areas in which base classifiers are fully trained. The final solutions and performance will be better as compared to integrated base classifiers and also the performance of base-classifier algorithms. Following are the few of the supervised techniques for classification involving both single classifier and ensemble classifier.

#### Support Vector Machine (SVM)

SVM is most widely used machine learning supervised learning technique, introduced by Vapnik (2000) [34]. Giveki *et al,* 2012 [35] modified cuckoo search for automatic diabetic diagnosis which was based on weighted SVM [1]. The Bhatia *et al*, 2006 [36] successfully classified heart disease by implementing a SVM-based classification system using integer coded genetic algorithm feature selection method which selected relevant feature from Cleveland Heart disease database. This maximized the accuracy of SVM classification by reducing the number of irrelevant features which could potentially hinder the performance of the typical SVM classifier to classify heart disease. On similar approach used SVM to classify heart failure patients [37].

*Method:* SVM basically develops separating hyper-planes to enhance the classifying margin between positive dataset and negative dataset. The nearest two points to the hyper-plane in a pair are called support vectors. A set of labelled input training data with positive and negative input samples are fed to SVM classifier from which it learns for linear decision boundary to be able to efficiently discriminate the unseen genes of experiment. A key attribute of SVM is that it can take only fixed length of the input vector.

First of all genes in training set and test set needs to be transformed into feature vectors, then training vectors are fed to SVM for constructing classifier. The output for SVM is a predicted class for each sample in the test set [38]. Similar approach is employed for RNA samples and other large datasets prone to high dimensionality problem. SVM first constructs a hyper-plane according to training RNA-Seq dataset, and then maps an input vector into higher dimensional vector space (also called Hillbert space). Then mapping is usually done via kernel function. Software based on LIBSVM algorithm was employed for SVM classification and regression [38].

#### Bagging support vector machines (bagSVM)

A single SVM classifier is incapable of learning the exact parameters that could be generalized to all datasets. Therefore, bagging ensemble of several SVM classifiers can sort this issue. A machine learning approach called Bagging support vector machines (bagSVM) is a bagged ensemble of SVMs for classification of RNA-Seq data. bagSVM trains each of the SVM separately using bootstrapping technique then combines result from each trained SVM model using majority voting [2].

*Method:* Zararsiz *et al,* 2014 applied a bagSVM on simulated data [2] and real dataset [2]. The simulated datasets taken here was generated using a negative binomial distribution model and two of the real RNA-Seq datasets taken were obtained from public resources. Then DESeq normalization was performed over each of them to reduce batch effects. Variance stabilizing transformation (vst) was performed. Further the genes are ranked in decreasing order of their significance. Since BagSVM is a bootstrap ensemble method. For a given test data several SVM classifiers are trained individually using learning algorithm through bootstrapping technique and then are summed up using aggregation technique. SVM ensemble is done through repeated random sampling with replacements of training dataset, **K** replicated training sets are generated. Each sample **x**_**i**_ may a repeatedly occur several time or might be not at all in any particular replicate training set TR. Each of the TR will be further used to train a specific SVM classifier [2]. If datasets has class labels for different classes, then these labels can be used to train a decision tree for classifying raw data [1].

#### Classification and regression trees (CART)

Binary tree classifiers can be constructed by iteratively splitting the data into two child subsets (nodes). Each of the terminal nodes is assigned a class label and the partition corresponds to classifier. The three rules for tree building are (i) selecting appropriate splitting rule; (ii) split-stopping rule to consider a node to be terminal or to continue splitting and (iii) assignment of each terminal node to specific class. CART was developed by Breiman *et al,* 1984 [39]. It is a popular decision tree classifiers implemented in several fields. It randomly chooses a split by applying “Gini indexing” technique that effectively reduces noises at each node.

Let the probability of class *i* at node *j* be *p*(*i* | *j*), then the calculated Gini index will be 1- *Σ_i_* (*p*2(*i* | *j*). The generated tree is gradually trimmed upward approaching reduced sequence of subtrees in order to grow a maximal tree. Then a cross validation method is implemented in order to identify the subtree with lowest error rate of classification. Further the class is chosen for each terminal node to minimize estimate of resubstitution for mis-classification probability [2, 39].

#### Random forests (RF)

RF is also an example for ensemble method. It is collection of several untouched CART trees. In RF, predictions from several weak classifier trees of forest are averaged to result in a strong classifier to give most accurate prediction result [40]. Firstly, the training set for individual trees is produced by bootstrapping the original sample data [2]. Then bagging of each of the decision trees is done after training it over this bootstrapped sample dataset. Then RF selects few features to split at each node in order to grow tree [8]. It is more flexible than other machine learning classification approaches like SVM, etc. It has a few tunable parameters.

Let for *k*^*th*^ tree, a random vector *Ɵ*_*k*_ is generated. It is unaffected by previous random vectors with same distribution as *Ɵ*_*k*_… … … *Ɵ*_*k-1*_. Using the training dataset and *Ɵ*_*k*_, a tree can be grown to construct a classifier *h(x, Ɵ_k_),* where *x* is an input vector. In bagging step, random vector *Ɵ* is generated from counts obtained by *N* training set.

*Definition:* A random forest is a ensemble classifier that consists of several tree-structured classifiers {*h*(*x, Ɵ_k_*), *k* = 1, …} where each *Ɵ*_*k*_ independent random vectors and each tree plays role in determining most appropriate class at input *x*. RFs gives better result than boosting and adaptive bagging without altering the training set. They are accurate and reduce biases [40].

When several enough trees are generated from the procedure, then there is need to find most appropriate class for prediction [41]. RF involves bootstrap resampling process (also called bagging) which is partitioning of a random feature. Here, at each tree node a subset of features is selected for modeling prior to modeling as in Fig. 9 [42]. Since bootstrapping is a sampling process by iterative replacement from the training data, some sequences may be missed from the sample and some may be repeated in sample. The missed sequences will be out-of-bag (OOB) sample. Usually, forest is generated using approx. 2/3 of training sequences and 1/3 is missed as OOB. Since OOB sequences are not involved in forest construction, so can be used in predicting performance. The 2/3 of training sequences is used to construct forest. The average error of 1/3 of OOB training data points is calculated [8]. Thus bootstrap resampling process leads to simultaneous cross-validation with help of OOB making OOB error rates a key for performance measurement for random forest algorithm for prediction [42]. Values of each feature are then further exchanged among rest of the 2/3 training data and again the OOB error is calculated with the help of new trained. In order to meet curse of dimensionality, Random Forest approach was applied to rank importance of these features based on some selected subset. Only top few features are then considered for further study [8].

**Fig. 9.**
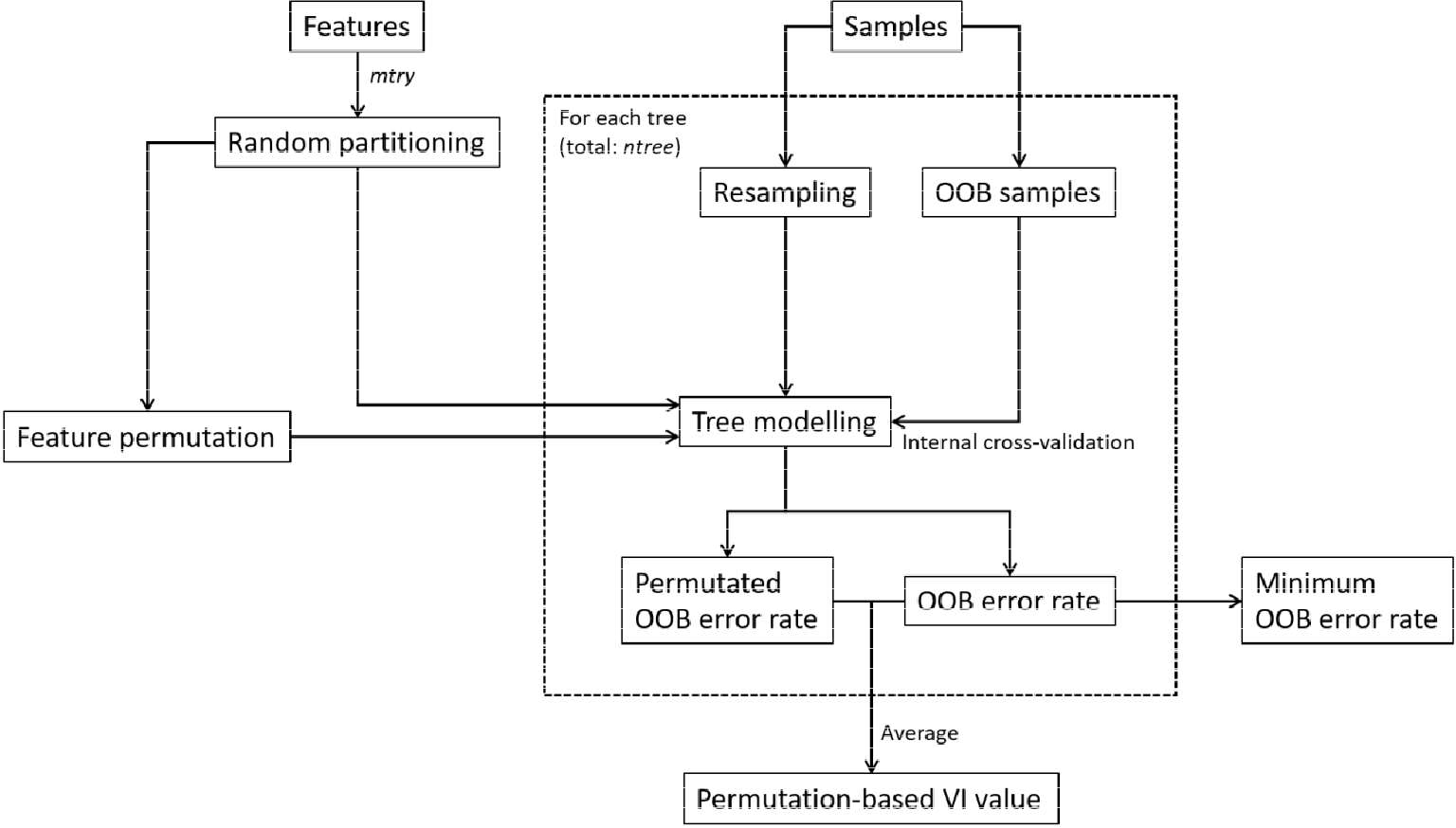
Flowchart of the core algorithm for random forest [42].

By partitioning features, the modeling biases of dataset or feature can be effectively reduced. In this step, in order to generate a decision tree, a non-parametric methods (CART) or parametric methods is applied to each of the randomly selected features. Hence, we get a collection of several decision trees as a result in form of Random Forest model with unbiased feature evaluation and greater classification capability [42].

*Method:* The Fig. 4 shows a flowchart of the random forest based feature selection classification method. The process outline is as follows:

- A univariate filtering is performed over validated miRNA-Seq dataset using analysis of variance (ANOVA) or Student’s t-test in order to eliminate statically insignificant.
- A bootstrapping method of Random Forest is performed on the filtered dataset to calculate the variable importance (VI) associated to each of the feature variables.
- Permutation-based VI value is assigned to all the features. This VI value is actually the average difference between error rates calculated from all the RF trees involved.
- VI values are used to rank all features in descending order.
- Then standard deviation (SD) for each of the feature along with their corresponding VI ranks is used in CART regression modeling to calculate the minimum SD.
- This SD is then used as threshold for the VI rank-based feature elimination step. The more relevant feature has larger variance in VI values.
- The remaining features then undergo further bootstrapping (50 times) process SFS-like process starting from feature ranked at the top of the VI ranking.
- Finally, the subset of features with mean OOB error rate less than the minimum OOB error rate and one standard deviation is considered to be the final result [42].

In case of identifying miRNA for a specific process, identification of maximum number of features is recommended since their regulatory responses are statistically correlated when establishing a RF feature selection procedure [43]. The recursive RF approaches shows maximum performance for correlated features so suitable for handling gene expression datasets [44]. It is also to be noted that evaluation of computational power and time requirement for this process is also necessary as we have already seen that Boruta though identified all the features but required large sample sizes and computational power [45].

#### Poisson linear discriminant analysis (PLDA)

Witten (2011) proposed a classification method for RNA-Seq data called Poisson linear discriminant analysis (PLDA) [22] which assumes that RNA-Seq data follow Poisson distribution. When biological replicates of RNA-Seq data are not available, Poisson distribution is suitable for modeling purpose but when it is available, it is not recommended use Poisson distribution due to overdispersion issue where variance of data exceeds its mean [24]. Overdispersion highly affects the performance and accuracy of classification [46].

PLDA may be a good choice for count based classifier after power transformation of data in all dispersion settings [47]. Recently, a few studies were performed to classify the sequencing data involving PLDA which concluded that PLDA classifier is somewhat similar to diagonal LDA and performs satisfactory over sequencing data [22, 48].

*Comparison between these classification models:* Zararsiz *et al*, 2014 [2] compared result of models with implementing classifiers single SVM, classification and regression trees (CART), Poisson linear discriminant analysis (PLDA) and random forests (RF), it was found that CART was not suitable for overdispersed data and others were well for it. Performance of PLDA was better than RF and single SVM, but when data was more overdispersed, bagSVM tend to be the best classifier [2]. The Fig. 10 represents the performance of classification methods on overdispersed dataset such as RNA-Seq data.

**Fig. 10.**
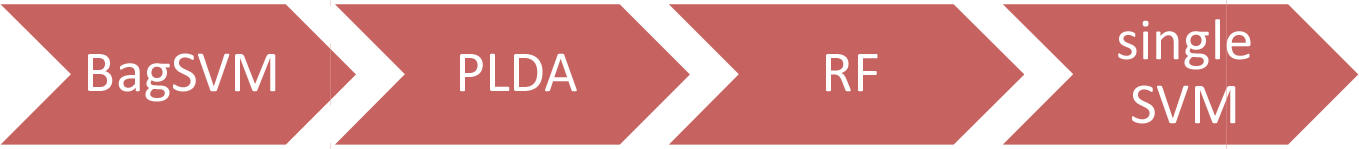
The performance order of classification method for overdispersed RNA-Seq data in decreasing order.

### 4.2 Unsupervised learning methods

Unsupervised learning process does not use class labels for learning algorithm. Clustering is the most widely used unsupervised technique that groups the objects into clusters such that the distance between objects of same cluster is minimum, and between different cluster is maximum [49, 50]. The objects of same cluster are similar in all features involved. The mutual distance between a pair of object is seen as the major challenge of clustering technique. This issue is met by formulating proximity measures like Cosine distance, Euclidean distance, City block distance. In clustering process while calculating the distance between pair of object, almost every feature is inferred [1].

In RNA-Seq dataset where several features are involved, researchers are interested in grouping similar objects over a feature subset [51]. Hence biclustering technique a variant of clustering emerged. Here each of the generated bicluster is formed by their respective feature subset. Like clustering, biclustering is also a two dimensional data analysis approach, where each of the features is corresponding object value of the object. While quantifying feature of an attribute each feature of the attribute is treated as object value of object [1]. Due to advances in data generation technologies it is now possible to track the dynamic nature of attributes of objects over multiple time instances. Hence three dimensional data analysis approach evolved leading to generation of another variant of clustering called triclustering. A tricluster thus generated consist of objects which are similar regarding subset of feature and also subset of time points [52]. It leads to a simultaneous grouping of objects with feature points and time points.

The clustering methods can be classified into partitional clustering, hierarchical clustering, density-based clustering, graph theoretic clustering, and soft computing-based clustering methods depending on their mode of working, as shown in Fig. 11.

**Fig. 11.**
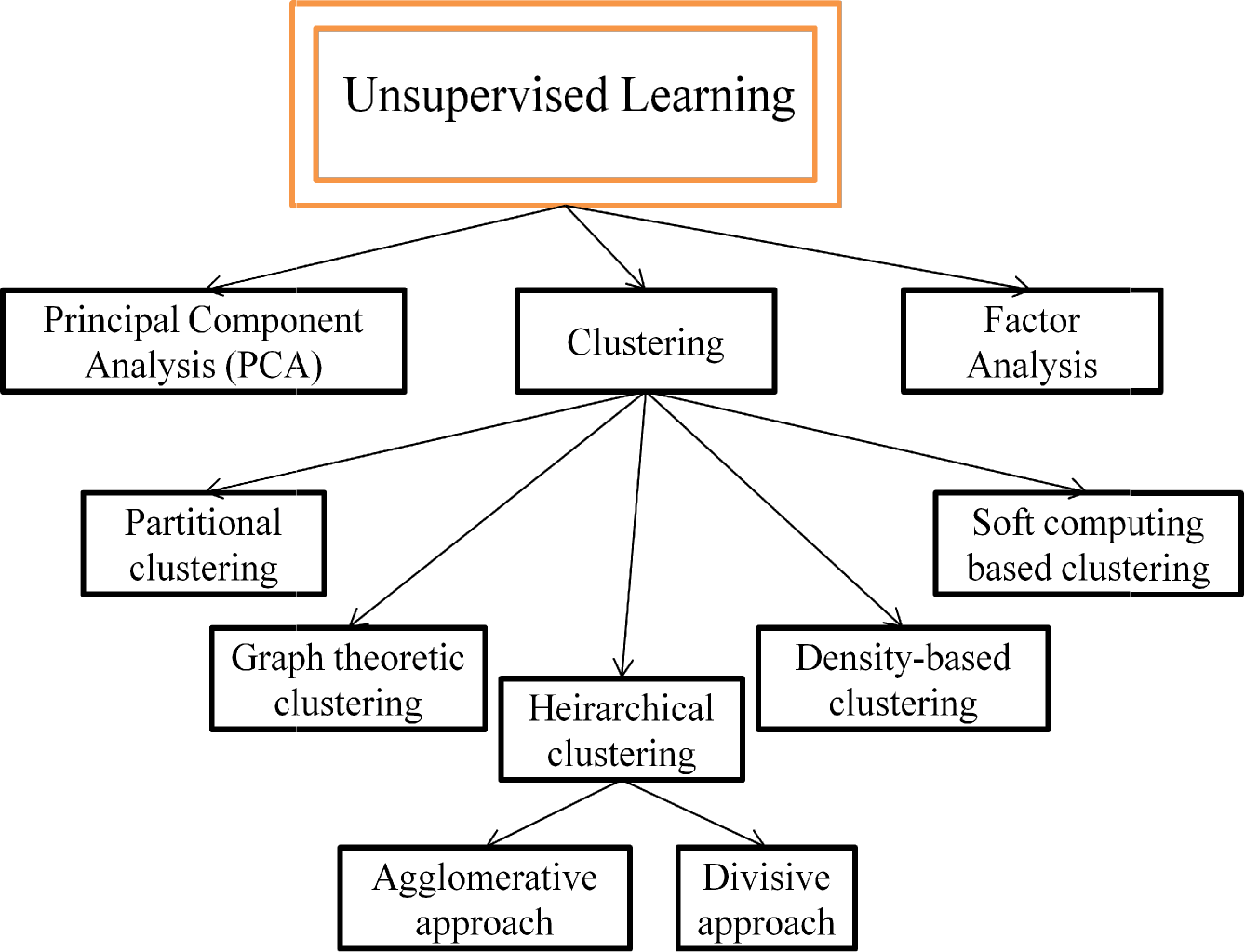
Taxonomy of unsupervised learning.

#### Partitional Clustering

This approach submits objects either of the k clusters formed. The k is actually a user defined parameter that optimizes the criteria of clustering. In this K means approach of clustering object is assigned iteratively to the nearest centroid of cluster until all objects get assigned.

CLARANS [54] is its implementation. CLARAN uses sampling during neighborhood search operation. It identifies the spatial structures in a test dataset. It developed two spatial data mining algorithms that can discover knowledge which was earlier difficult to find with existing spatial data mining algorithms by establishing relationships between spatial and non-spatial attributes [54].

#### Hierarchical clustering

This method is subdivided into agglomerative and divisive hierarchical clustering methods. In agglomerative clustering, individual object is considered a separate leaf node which is gradually clustered in bottom up approach of the tree iteratively to reach its root [1].In divisive approach, the large cluster (root) having all the objects (nodes) are iteratively redefined and split into daughter nodes in order to reach leaf node ultimately [53].

Its implementation is BIRCH (Balanced Iterative Reducing using Cluster Hierarchies) for agglomerative [64], CURE for divisive [65].

#### Density-based clustering

This method relies on measuring the density of node using neighborhood approach and then categorizing cluster by dense area boundaries by lower dense [55].

Its implementation is DBSCAN [66]. It is a density-based clustering method that starts from an initial object and includes objects from its neighborhood iteratively if they satisfy a user defined threshold to form a cluster. DENCLUE [67] uses kernel density function to find a local maximum so as to call it a cluster [1].

#### Graph theoretic clustering

This method uses the basic properties and concepts of a typical graph theory.

Its implementation is “Chameleon”. Chameleon [68] is a graph theoretic clustering method that uses k-nearest neighbor graph. Here, edges get deleted iteratively deleted if associated nodes are not included in the k-nearest neighbor sets [1, 56].

#### Soft computing-based clustering

This method makes use of fuzzy sets, neural networks and other soft computation tools. One of its examples is fuzzy c-means. It is a crisp clustering method that allows an object to be associated with several cluster on a rule that sum of the membership feature of object to associated SOM consists of vectors with high dimensionality in a 2-D space. It iteratively allows objects to join dense region cluster [1, 57].

### 4.3 Hybrid Method (Deep Learning)

The increase in dimensionality of biological data is a great challenge to conventional analysis techniques like simple Supervised (Naïve Bayes, ANNs, Random Forests, etc.) and Unsupervised (Clustering, etc). Artificial neural networks (ANNs) is a powerful approach for learning complex patterns at multiple layers inspired from biological neurons responsive to brain processing. ANNs implement learning by involving some sorts of simpler base elements arranged in complex networks. However, their internal representations are difficult to be interpreted and training these deeply layered models has been algorithmically challenging and is statistically prone to overfitting [69]. Therefore, the modern machine learning methods like Deep Learning technique processes large datasets such as RNA-Seq data to find the hidden structures within them enabling more accurate predictions [63]. Deep learning is actually a modified multilayered ANNs [69]. Deep learning is models higher level data abstractions using some model architectures composed of several non-linear transformations. Deep learning is an efficient learning method which involves deep architecture to perform some intellectual learning (Eg. learning the features). The deep architecture is the multilayer network with interconnected adjacent layered module.

Depending on the learning nature of these layered modules, the existing deep learning algorithms can be classified into as follows (Fig. 12 and Fig.13):

- *Generative* (e.g. deep auto-encoder, deep Boltzmann machine and deep Belief networks (DBNs)),
- *Discriminate* (e.g. convolutional neural networks (CNNs) and deep stacking networks (DSNs))
- *Hybrid architectures* (e.g. deep neural networks (DNNs)) [14]

DNNs require stochastic gradient descent which makes simultaneous network parameter learning virtually impossible. Therefore, the deep stacking networks (DSNs) were introduced. The basic DSN architecture consists of several stacking modules which are individually a shallow multilayer perception. It uses convex optimization for learning the perceptron weights. “Stacking” is achieved by concatenating the output predictions of all previous modules, original input vector form new “input” vector in the new module. Deep learning architectures are convolutional CNNs, DBNs, DNNs and DSNs which have been successfully shown better results in computer vision, automatic speech recognition, natural language processing, and music/audio signal recognition, etc [14].

**Fig. 12.**
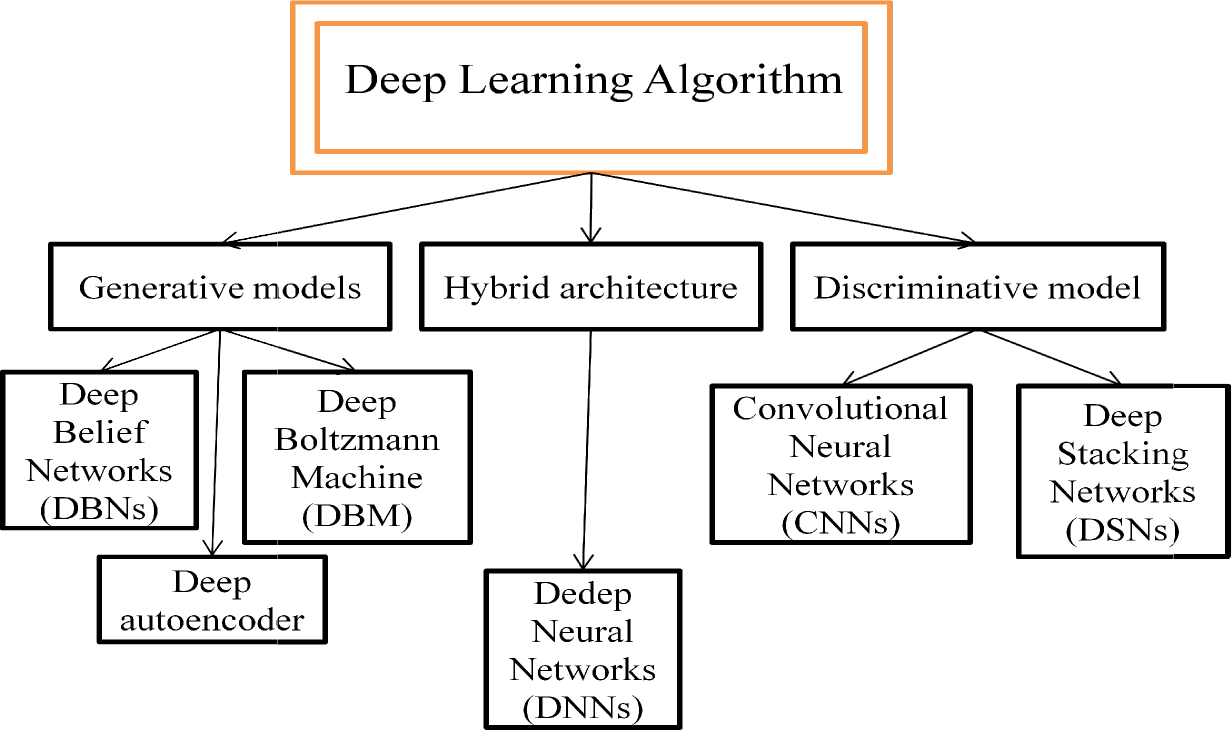
Deep learning classification.

**Fig. 13.**
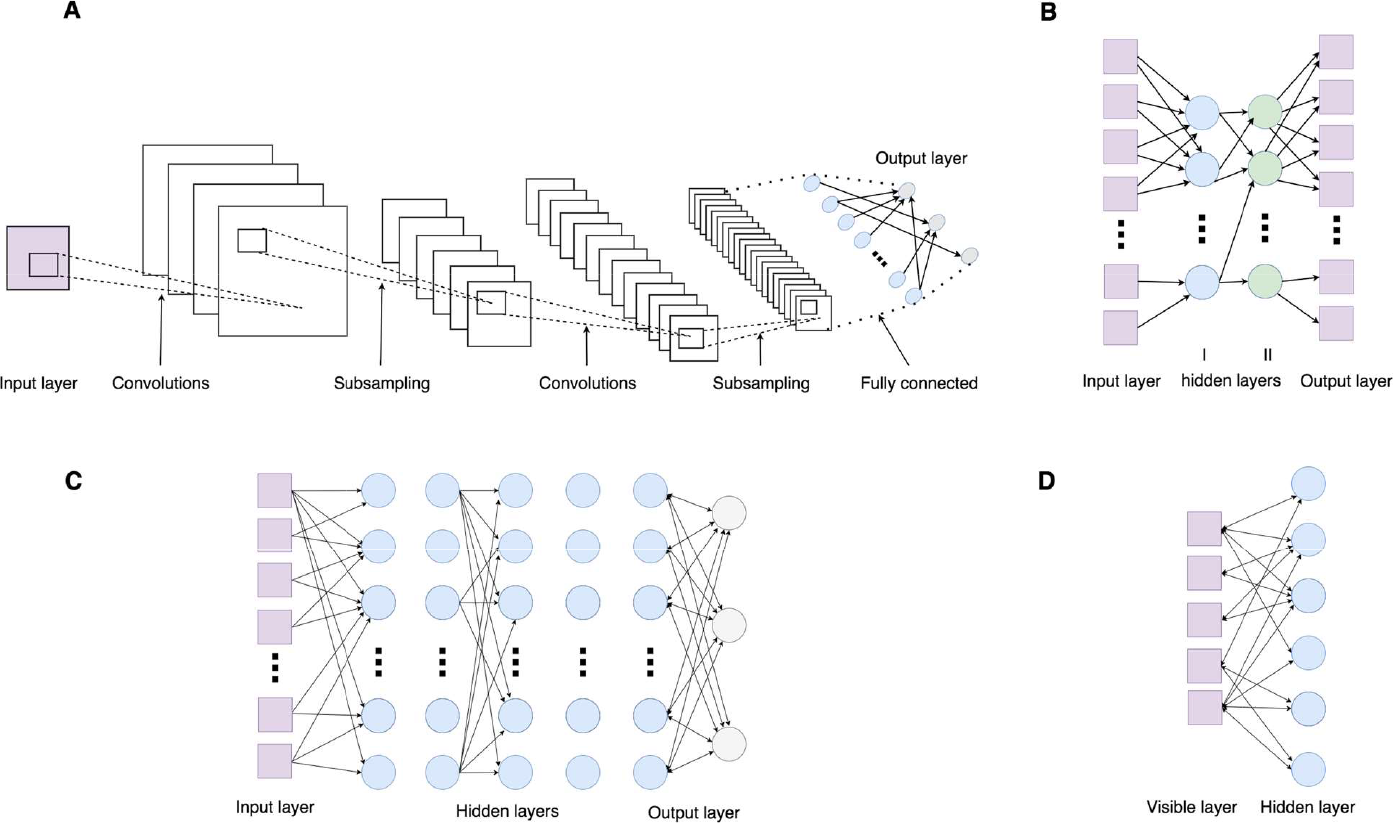
Four of the most popular classes of deep learning architectures in biological data analysis. (A) Convolutional neural network (CNN): has several levels of convolutional and subsampling layers optionally followed by fully connected layers with deep architecture. (B) Stacked autoencoder: consists of multiple sparse autoencoders. (C) Deep belief network (DBN): trained layer-wise by freezing previous layer’s weights and feeding the output to the next layer. (D) Restricted Boltzmann machine architecture: includes one visible layer and one layer of hidden units [7].

These approaches have also been applied to several computational biology problems dealing with regulatory genomics and image analysis [63].

Most popular approach to deal with high dimensional data such as RNA-Seq data involves DNNs. DNNs are able to handle large datasets with high dimensionality, sparse, noisy data with nonlinear relationships such as in case of transcriptomics and other -omics data in biology. DNNs have high generalization advantages. Once trained on a dataset, it can be applied to other large datasets. Therefore, it can interpret heterogeneous multiplatform data efficiently such as Gene expression datasets reducing the issues like dimension reduction and selectivity/invariance [7].

Deep learning algorithms such as DNNs can be used in biological research field like in annotation, biomarker development, drug development and discovery, transcriptomic data analysis, etc. A combination deep learning with shallow learning methods like supervised, unsupervised, and reinforcement learning can be applied to biomedical datasets such as RNA-Seq data to understand the biological mechanism for disease and help in developing personalized medicine.

Fan & Zhang, 2015 developed lncRNA-MFDL [14] predictor to identify lncRNA by fusing several features (k-mer, the secondary structure and the most-like coding domain sequence) together and using deep learning classification algorithm [14]. In one deep learning application using five tissue-specific RNA-seq data sets, a DNN was developed using hidden variables for features in both genomic sequences and tissue types and was shown to outperform Bayesian methods in predicting tissue splicing within individuals and across tissues, specifically the change of the percentage of transcripts with an exon spliced (PSI), a metric for splicing code [70]. Features from gene expression were extracted with regions of noncoding transcripts (miRNA) using DBNs and active learning. Here deep learning feature extractors were used to reduce the dimensionality of six cancer data sets and outperformed basic feature selection methods [71].

#### Important considerations for implementing deep learning

One of the challenges faced while implementing Deep learning in solving a problem is selecting the appropriate DNN type for the task. Few of the following consideration should be kept in mind while developing DNNs for a particular application. DNNs can be classified into following three major categories [7].

*Networks for Unsupervised learning:* They ensure data correlation by identifying combined statistical distributions with some particular kind of associated classes when available. Bayes rule can further be implemented to lat discriminative learning machine classification.

*Networks for Supervised learning:* They provide maximum discriminative power in classification. They are trained only with labeled data where all outputs must be tagged.

*Hybrid or Semi-supervised networks:* It is used to classify data using the outputs from generative (unsupervised) model. The data is used to train the network weights prior to supervision to speed up the learning process prior to the supervision stage.

#### Methodology for deep learning in context of computational biology

Angermueller *et al.* (2016) gave a systematic method to implement deep learning method for biological dataset classification (Ex. RNA-Seq data). These steps are enlisted below [63].

##### Data Preparation

The more informative features of training dataset usually result in better performance. Therefore effort should be spent on collecting, labeling, cleaning and normalizing data. Machine learning models need to be trained, selected and tested on independent data sets to avoid overfitting and assure that the model will generalize to unseen data.

###### Dataset size

For a supervised learning setting sufficient labeled training samples should be available to fit complex models. The number of training samples should be higher to avoid overfitting though special architectures and model regularization are present to deal it training data are scarce.

###### Partitioning data into training, validation and test sets

For DNNs implementation initial task is to partition r dataset (e.g. RNA-Seq data) into training, validation and test sets. The training set is then subjected to learning process to learn models with different hyper-parameters. Then they are assessed validation set. The model with best performance (prediction accuracy) is selected and further evaluated on the test set to quantify the performance on raw data or for comparing to other methods. If the data set is small in size, k-fold cross-validation or bootstrapping can be used for evaluation.

###### Normalization of raw data

When the features of RNA-Seq data are skewed (biased) due to some batch effects, log transformation or similar normalization methods can be adopted to accelerate the training and identification phase in “Classification” process.

##### Model Building

###### Choice of model architecture

The default architecture is a feedforward neural network with several fully connected hidden layers. The convolutional architectures are suitable for multi and high-dimensional data. Recurrent neural networks can capture long-range dependencies in sequential data of varying lengths, such as text, protein or DNA sequences.

###### Determining the number of neurons in a network

The optimal number of hidden layers and hidden units is problem dependent and should be optimized on a validation set without overfitting the data. Higher the number of hidden layers and unit, higher is the number of representable functions.

##### Model Training

It is performed to detect the parameters that would minimize the objective function which measures the fit between predictions from the parameterized model and actual observation. Stochastic models are used to train the deep models.

Some of the central parameters for ANNs are *learning rate, batch size, momentum rate, weight initialization, per-parameter adaptive learning rate methods, batch normalization, learning rate decay, activation function, dropout rate.*

*Parameter initialization:* Usually model parameters are anlysed randomly. These can be sampled independently from normal distribution of small variances or variance scaled inversely by hidden units in input layer.

*Analysis of the learning curve:* This is done in order to validate the learning process done earlier. If the curve decreases slowly, learning rate is small and needs to be increased. If curve decreases steeply, learning rate may be high. High learning rate leads to fluctuating learning curves.

*Training and validation performance monitoring:* Along with monitoring the loss in training it is also necessary to monitor target performance like accuracy of training and validation sets. Low validation performance compared to training performance signifies overfitting of data. Overfitting results from too complex model relative to the size of training set. Thus by decreasing the model complexity (decreasing number of hidden layers and units) or by data augmentation (i.e. increasing size of training dataset), overfitting can be avoided.

#### The basic learning algorithm for Deep Stacking Network

The general algorithm of Deep Stacking Network (DSN) consists of the weight parameter called input network weight matrices **W** and the output network weight matrices **U** in each module is learnt from training dataset with the help of basic tuning algorithm and learning algorithm.

Let there be training vectors **X = [x_1_, …, x_i_, …, x_N_]** and some target vectors **T = [t_i_, …, t_i_, … t_N_]** for total **N** number of training samples. The output of this DSN learning module will be **y_i_ = U^T^ h_i_**; where **h_i_ = σ(W^T^ x_i_)** is hidden layer output and **σ (.)** represents the sigmoid function. The loss function error of the mean square error is utilized to learn the output weight matrices assuming that the lower layer weight matrix **w** is already given. This is achieved by minimizing the average of total square error **E = ||Y-T||^2^ = Tr[(Y-T)(Y-T)^T^]** to learn the parameter. If **w** is fixed, the hidden layer values **H** can be calculated and thus the upper layer weight matrix **U** in each module can be determined by calculating the gradient 
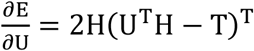
to **0.** This result in equation, 
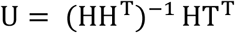

It is to be noted that in DNS module, all the weight matrices **w** should be set empirically. This can be done either by using various distributions for generating random numerals to set **w;** or by training Restricted Botzmann Machine (RBM) separately using contrastive divergence, then this trained RBM set **w.** The weight matrix **w** of DSN in each module can be further learnt by applying gradient descent as,
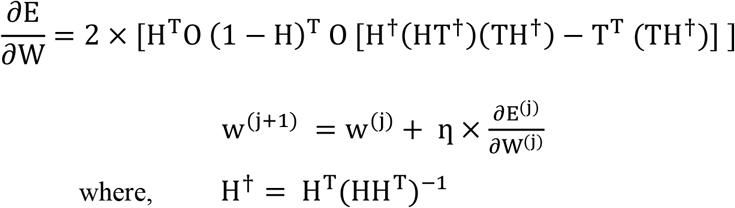

The symbol **O** represents the element-wise matrix multiplication, and **η** is the learning rate of updating the weight matrices **w** [14].

## 5 Tools and pipelines implementing machine learning approaches for RNA transcript identification and classification

Over the years, several tools and pipelines have been developed to find solution for bioinformatical problem. Tools developed prior to Big Data era were usually standalone. During last decade several data analysis tools and pipelines have been developed for identification and classification of RNA transcripts, coexpressed genes analysis of gene-gene network, etc.

### 5.1 miRNA-dis

It is used for identification of microRNA precursor. miRNA-dis uses SVM in order to predict structure order information. On the basis of distance pair, the feature vector can be constructed. It employed SVM. The SVM analyses the data and recognizes pattern for classification and regression analysis. SVM training algorithm builds a classifier model that assigns new test sample either of the categories. Hence it is a non-probabilistic binary linear classifier. In miRNA-dis LIBSVM algorithm [58] was employed which is based on SVM [38].

### 5.2 iSeeRNAs

It uses SVM model to detect lncRNAs by implementing several features [12]. iSeeRNA uses SVM model to identify the lincRNAs by using several features such as conservation, ORF, di- and tri-nucleotide sequence frequencies [14].

### 5.3 PredcircRNA

It is a machine learning approach which identifies circularRNA from lncRNAs by implementing multiple kernel learning. Firstly, different discriminative features like graph features, sequence compositions, etc are extracted. Then fusing these feature using multiple kernel learning. This analysis was based on Random Forest method [8].

### 5.4 MiPred

It is based on Random Forest prediction model. It differentiates real pre-miRNAs from pseudopre-miRNAs by involving a hybrid feature system with participating features such as structure-sequence composition, minimum free energy (MFE) of secondary structure and P-value of randomization. RF method implemented here showed 10% higher accuracy than triple SVM classifier because of ensemble of features [41].

### 5.5 rxDTree

It is used to estimate a fast and distributable decision tree on big data. It is implemented in classification and regression problems. It builds a decision tree in Breadth-first manner by constructing histogram for generating distribution functions of data [59].

### 5.6 CAMUR models

The Classifier with Alternative and MUltiple Rule-based (CAMUR) model includes some knowledge databases and a query tool. It extracts out several similar classification models. For this, it first computes classification model based on single rule, and then it calculates the power set of genes in rules. Then it removes these combinations from data iteratively if stopping criteria is not satisfied. Then again performs classification process until stopping criteria is verified [60].

### 5.7 QUBIC algorithm

The QUalitative BIClustering (QUBIC) algorithm uses qualitative or semi quantitative methods of gene expression data along with optimizing technique to solve biclustering problems within a short time period. It can efficiently recognize statistically significant biclusters [61].

## 6 Discussion and future challenges

Various computational tools have been developed for studying pathogen or virus from RNA Seq data by classifying them according to the attributes in several pre-defined classes, but still statistical tools and approaches to analyze complex datasets are still lacking. Moreover there is no ‘one fit to all’ technique for classification and analysis of RNA-Seq data therefore it’s a challenging task. Few of the challenges in RNA-Seq data classification in these contexts is enumerated as follows:

### Imbalanced data problem

The imbalanced data occurs in medical diagnosis along with other practical fields like banking, oil exploration, information retrieval, text classification, etc. This problem of imbalance is largely seen in miRNA-Seq data. It consists of both positive and negative example set. Only positive examples are required for validation in a biological experiment. Overrepresentation of these negative examples is very common. These can be filtered out but is not a solution for this overrepresentation and imbalance problem. Traditional machine learning methods of classification gives inferior performance for it. Therefore, it is highly recommended to develop efficient machine learning algorithms for classification of unbalanced data positive and negative resources in bioinformatics [4].

### Performance affected due to high density data generated

The real time analytics of RNA-Seq data had become harder to be processed due to speedy data generation. This is attributed to next generation and third generation sequencing techniques. Though batch mode of processing is applied to this high density data using distributed and parallel computing but still analytic performance get affected due to this big data [1]. Moreover bioinformatical data is massive and holds dimensionality problem while the number of instances is geographically distributed worldwide.

### Heterogeneous nature of data

The bioinformatical data generated today specially RNA-Seq data are heterogeneous in nature with several features sources summed in a single test sample. The traditional databases those are inferred for processing are based on a definite schema are unable to handle this heterogeneous data [1]. Since bioinformatical data are relatively heterogeneous in nature, they require several heterogeneous and independent databases for analytics and validation [62].

### Data is not in a uniform format

Since the big data from RNA-Seq is obtained from different sources and are widely heterogeneous in nature, they are obtained in different format. This increases the complexity of data analysis process. Therefore, more intelligent machine learning algorithms are required to classify this discrete formatted data [1]. Bioinformatical data are usually generated by many several independent organizations worldwide so there are high chances that same types of data may be represented in different forms at source. This results in need of highly sensitive analytics approach [62].

### Challenges posed in training deep learning models

Training deep learning models poses far greater challenges than training shallow models, both for defining model parameters and model structures. Deep learning models are still not optimized, lack an adequate formulation, require more research, and rely heavily on computational experimentation.

### Deep learning is not suitable for sparse dataset

There is no single method which can be applicable to all sorts of problems. The choice of deep learning approach and usage totally depends on nature of problem dataset. Conventional analysis approach is no doubt still advantageous when dataset is sparse and need to be just statistically considered. This is still not suitable in deep learning.

### The “Black Box ”

They learn by simple association and co-occurrence and have limited sources from which could interpret internal representations. The high dimensional biological data is not easily interpretable and needs additional quality control and pre-processing. Thus, DNNs lack transparency and interpretability concerning to structural relationships common in biology without human input.

### Need for large data sets to avoid overfitting

It is not suitable for sparse dataset and require large training dataset which might not be always available specially the RNA-Seq data. When data sets is not sufficiently large, problem of overfitting can be encountered where training error is low and test error is high which deteriorates the generalization property of model and reduces the predictive performance of model. Overfitting can be regularized but still unbiased and noisy biological dataset such as RNA-Seq data may face overfitting.

### The computation costs

Finally, while shallow learning require few computational resources, but deep learning are usually computationally intensive and time-consuming and often requires access to and programming knowledge for graphics processing units (GPU) [7] and multi-processing distributed architecture over cloud.

These mentioned challenges paves the path for research in developing more efficient tools and technologies for Big RNA-Seq data analytics in field of bioinformatics.

## 7 Conclusion

The transcriptomic analysis involves quantifying variation in various types of transcripts like mRNA), long, lncRNA), miRNA), etc. to gather wide range of information from data such as biomarkers for disease, etc. The Gene Expression data can be represented in form of Gene Expression Matrices which holds the quantitative value of expression. These obtained data faces curse of dimensionality problem signal-to-noise in the dataset. Thus, statistical analysis for the dataset becomes difficult. This high dimensional data can be handled either by dimensionality reduction through feature extraction methods (SVM or Random Forest algorithms) or by pathway analysis; or by using such methods which are less sensitive to high-dimensional data such as Random Forest or deep belief networks. One can use either deep learning or shallow learning for classification of dataset, depending upon the type of raw data. Shallow learning such as SVMs, ensemble methods like bagSVM, Random forest is used suitable for sparse dataset but not the deep learning as it may encounter overfitting problem. But deep learnt networks can effectively discover high level features thus improving the performance over traditional shallow learning models, thus they can benefit researcher with additional information about biological datasets such as RNA-Seq data.

We can conclude that deep learning is a better choice for classification of large RNA-Seq data for biomedical investigations. But due to unavailability of large enough dataset, this approach may encounter overfitting problem. Moreover, this approach is still costly. Thus, there is need of development of further classification models which could successfully overcome the pitfalls of deep learning as well as shallow learning without loss of any important feature from dataset.

